# Immunogenetic diversity and haemosporidian parasitization in European bluethroats (*Luscinia svecica*): do diverse genes warrant fewer parasites?

**DOI:** 10.1101/2025.09.03.673962

**Authors:** Dragomir Damnjanović, Masoud Nazarizadeh, Václav Pavel, Bohumír Chutný, Arild Johnsen, Milena Nováková, Jan Štefka

## Abstract

The health and fitness of vertebrates are constantly challenged by environmental stresses, which include exposure to diseases that pose significant evolutionary pressures on immune genes. The bluethroat (*Luscinia svecica*), has been extensively studied for its haemosporidian parasite diversity across Europe showing a high diversity and prevalence rate. Given possible evolutionary pressure exerted by these pathogens, the present study used high-throughput amplicon sequencing to explore the genetic diversity and selection mechanisms of three immunity genes (MHC I exon 3, TLR3 and TLR4) in three European bluethroat populations. Specifically, we seek to (1) characterize the genetic diversity of these immunity genes, (2) detect selection signatures shaping their diversity, and (3) investigate the relationship between these immunity genes and haemosporidian parasites (*Plasmodium* and *Leucocytozoon*). Selection analysis was conducted using *FUBAR* and *SLAC* methods, whereas GLM regression was employed to explore the correlation between MHC genes and haemosporidian parasites. Despite geographical and ecological differences, nucleotide and haplotype diversities were similar across all populations. Most frequent haplotypes were found to be shared by both red-spotted (*L. s. svecica*) and white-spotted (*L. s. cyanecula*) bluethroat subspecies. The “insular” Krkonoše population, despite being biogeographically peripheral and experiencing minor inbreeding, did not show significantly reduced immunogenetic diversity. Selection analysis revealed a higher presence of purifying selection in TLR genes and a combination of purifying and diversifying selection in the MHC gene, reflecting their evolutionary constraints and functional importance, while haemosporidian parasite pressure was not a major driver of genetic diversity.

## Introduction

The integrity of individual health and fitness in vertebrates is constantly challenged by a wide range of environmental stresses. Infectious diseases caused by various pathogens of viral, bacterial and parasitic origin represent one of the major evolutionary pressures that drive the diversity of vertebrate immune genes (Gokhale et al. 2013; Tellier and Brown 2007). Final infection outcome is influenced by the synchronized response of the genes that alter both the innate and adaptive immune system. Mechanisms of adaptive immunity operate through executing coordinated reactions to environmental antigens which are mediated by the major histocompatibility complex (MHC) genes. These highly polymorphic families of genes are characterized by the ability to produce vast diversity of polypeptide molecules that target both intra- and extracellular antigens. Located on cellular surfaces, the polypeptide binding regions (PBRs) possess an affinity for specific antigenic motifs (Murphy 2012; Robinson et al. 2003; Tizard 1992). Therefore, the adherence of antigen fragments to the PBR region modulates the behavior of antigen-specific lymphocytes that accurately dispense a finely tuned immunological answer coordinated with CD8-bearing cytotoxic T lymphocytes and antigen-presenting cells (Boyd et al. 2012; Murphy 2012; Tizard 1992). Involvement of antigen-presenting cells enables memorization of encountered antigen polypeptide motifs for the future interactions with the same type of antigen and ensures increased protection against repetitive invasion with the same antigen (Kasahara et al. 1995; Murphy 2012; Tizard 1992). In contrast to the adaptive immune system, the innate immune system provides an immediate answer through secretion of soluble polypeptides and enzymes that directs effector cells toward the initial microbial breach. Stimulated phagocytic effector lymphocytes (i.e. macrophages, granulocytes and dendritic cells) pursue pathogens that bypass respiratory and digestive mucosa. The extracellular domain recognizes a wide range of pathogen-associated molecular patterns (PAMP) motifs that execute the initial series of the nonspecific immune response. The behavior of phagocytic cells is modulated by transmembrane TLR domains that are capable of distinguishing antigen components by recognizing PAMPs molecule motifs characteristic for microbes (Murphy 2012; Netea et al. 2012).

Given that pathogens pose an evolutionary pressure on avian populations, the antagonistic co-evolution of immunity genes and respective parasite fauna is an everlasting process (Garamszegi and Nunn 2011; Hill 1991; Hill et al. 1991). Presumably, classic hypotheses postulate that high MHC genetic diversity is shaped by various models of balancing selection (*heterozygosity advantage, rare-allele advantage* or *fluctuating selection*) through continuous interactions between the parasite pressure and immunity genes (Edwards and Hedrick 1998; Takahata and Nei 1990). Consequently, continuous parasite selection will maintain sufficient genetic diversity at the inter-population and even inter-species level across the geographical and temporal scale (Spurgin and Richardson 2010). The overall efficacy of the immune response, mediated by the MHC genes, is augmented by high allelic heterozygosity that provides a conduction of wider array of pathogen motifs. Simultaneously, other mutually non-exclusive patterns of balancing selection (i.e. *fluctuating selection* or *rare allele advantage* selection) can maintain optimal MHC genetic diversity. Furthermore, disassortative mating is expected to elevate MHC diversity (Edwards and Hedrick 1998; Hughes and Hughes 1995; Penn et al. 2002; Sommer 2005). In contrast to MHC genes, TLR receptors show limited haplotype variation on individual and population level, as the evolution of the TLR genes is shaped predominantly by purifying selection manifested through the removal of deleterious non-synonymous mutations (Chapman et al. 2016; Mukherjee et al. 2009). Strong presence of purifying selection that maintains nucleotide diversity of TLR genes is reflected in the conserved molecular structure of TLRs across multiple vertebrate and invertebrate taxa. This highly restrictive endowment towards novel polymorphism is dominant as even a single amino acid mutation can diminish the efficacy of TLR’s PAMP-binding region (Keestra et al. 2008; Resman et al. 2009).

Detrimental health consequences inflicted by obligatory single-cellular haemosporidian parasites (Phylum: Apicomplexa) lead to anemia through the reduction of viable erythrocytes, which ultimately diminishes individuals’ mobility and resilience to environmental challenges. Thus, these vector-borne haemosporidian protists pose a strong selective pressure on (Davidar and Morton 2006; S. Knowles et al. 2010; S. C. L. Knowles et al. 2011; Marzal et al. 2005; Westerdahl et al. 2012). The bluethroat *(Luscinia svecica*) has been a focal non-model passerine species for several malaria studies, in which rich biodiversity of haemosporidian fauna has been described (Damnjanović et al. 2025; Hellgren 2005; Rojo et al. 2014; Svoboda et al. 2015), with an extensive volume of haemosporidian lineages deposited in MalAvi online database (http://130.235.244.92/Malavi/) (Bensch et al. 2009).

Certain MHC alleles have been reported to express increased sensibility to specific haemosporidian lineages represented by correlations between a presence or absence of particular MHC class I alleles or supertypes with the specific haemosporidian lineage (i.e. *Plasmodium, Haemoproteus* or *Leucocytozoon*) and reduced parasitemia in avian malaria infections (Loiseau et al. 2008, 2011; Westerdahl et al. 2012, 2013). Similarly, in this study we aim to explore a connection between the immunity genes (TLR3 exon 4, TLR4 exon 3 and MHC Class I exon 3) and the diversity and prevalence of haemosporidian parasites (*Plasmodium* and *Leucocytozoon*) in three European bluethroat populations. Specifically, we: (1) characterize the genetic diversity of immunity genes (TLR3, TLR4 and MHC class I exon 3 gene) in the bluethroat populations, particularly with respect to the Krkonoše population, which is close to extinction (Miles and Formánek, 1989, Chutný, 1991, personal observation); (2) Detect signatures of the selection shaping these immunity genes. (3) Explore relationship between immunogenetic diversity and infection by haemosporidian groups of parasites (*Plasmodium* and *Leucocytozoon*) using general linear regression models.

## Material and Methods

### Fieldwork blood collection and amplicon gene library preparation

Bluethroat blood samples were obtained from the three European bluethroat populations (Fig. 1). For the red-spotted bluethroat subspecies (*Luscinia svecica svecica*), 150 samples originating from the Krkonoše Mountains in the Czech Republic (50.688949°, 15.657069°) and 24 samples from the Abisko, Sweden (68.417089°, 19.207025°) were collected. The third studied population consisted of 24 white-spotted bluethroat subspecies (*Luscinia svecica cyanecula*) samples obtained in the Třeboň area (49.004196°, 14.772123°) in the Czech Republic. All samples were collected during multiple breeding seasons by Svoboda et al. (2015). Blood samples were drawn by sodium-heparinized microcapillary (Marienfeld, Germany) in volume of 10-50 uL and stored in Queen’s lysis buffer. DNA was isolated using DNeasy Blood and Tissue Kit (Qiagen, Germany) and stored in the 75 ml of EA buffer. MHC I class exon 3 gene amplification were performed using the *HN34/HN45* primer pair as in (Lyons et al. 2015). The amplification of TLR3 exon 4 and TLR4 exon 3 gene was performed using the *avTLR3F/avTLR3R* and *avTLR4F/avTLR4R* primer pairs respectively (Alcaide and Edwards 2011). PCR products stained with GelRed were visualized on 1.5% agarose gel at 90V in a 50x TAE buffer. Quality and optimization of preliminary gene sequences was assessed in Geneious Prime® 2020.1.2 software (https://www.geneious.com). Next generation sequencing (NGS) amplicon libraries were sequenced on Illumina Novaseq platform (150 bp pair-end reads) by Novogene (UK).

**Fig. 1.**
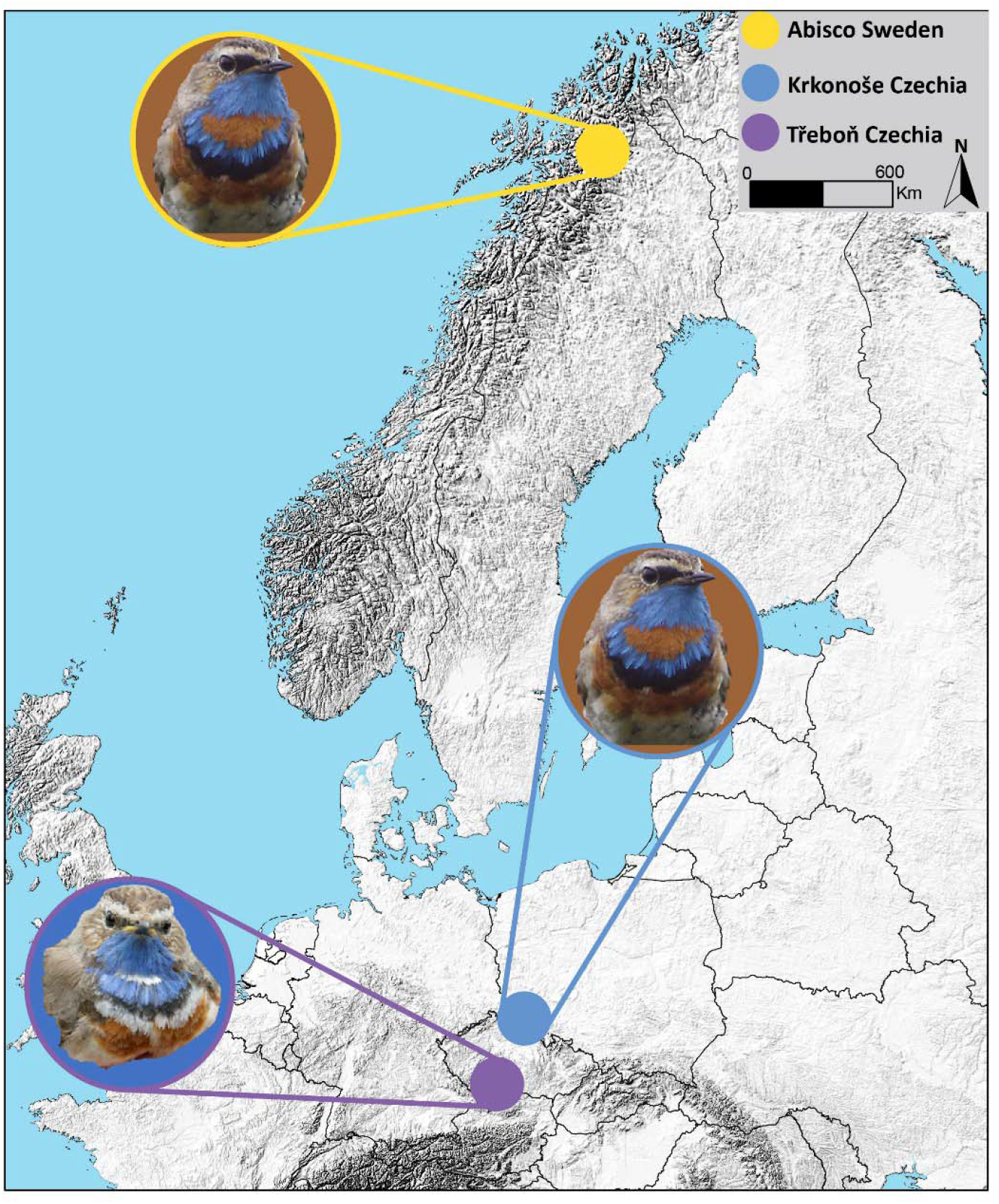
Map of studied bluethroat locations. Red-spotted bluethroat populations are found in Abisko (Sweden) and Krkonoše (Czech Republic). The southernmost population (Třeboň) represents the white-spotted bluethroat subspecies.

### NGS Amplicon library processing

Quality of raw amplicon sequences was analyzed in FastQC (phred score >30). Primers and barcode sequences and trimmed by Trimmomatic v0.39 (Bolger et al. 2014; Brown et al. 2017). Further optimization of the amplicon reads was processed using DADA2 R package 1.26.0 (Callahan et al. 2016). Both forward and reverse reads were filtered by the following *filterAndTrim* function settings (maxN=0, maxEE=c(2,2), truncQ=5, rm.phix=TRUE, matchIDs=TRUE, multithread=TRUE). Amplicon reads were dereplicated and merged using default DADA2 settings. Chimeric reads were removed according to the manual instructions. Two filtering criteria were set to filter the amplicon reads into the optimized dataset: 1) Haplotype variants that occurred at minimum 5% frequency per two sub-libraries were retained for further analysis; 2) haplotypes that contained less than 100 reads per individual were discarded (Gruenberg et al. 2019). The base pair quality, sequence length and translation reading frames of recovered sequences was evaluated using Geneious Prime V 2020.1.2 (Biomatters, Auckland, New Zealand). Sequences that contained irregular gaps and intersecting stop codons within all translation reading frames were removed from the dataset. Sequence length varied from 342 to 355 base pairs, whereas most alleles were 342 bp long. A total of 428 putative pseudogene and chimera sequences were removed from the dataset, due to insertions of 2 and 4 bp were removed from the dataset due to shifts in the reading frames. All removed pseudogene haplotypes were present in more than one occurrence. The MHC class I exon 3 dataset obtained from 200 bluethroat individuals contained 1179 sequences which was composed of 172 unique haplotypes (342bp). The dataset for the TLR3 comprised of 326 sequences, whereas TLR4 genes were composed of 435 sequences. A collection of 18 TLR3 and 69 TLR4 unique haplotypes were translated into 10 and 17 unique amino acid sequences, respectively.

### Gene polymorphism and diversity

Assessment of the immunogenetic diversity of the populations, statistics such as the number of unique haplotypes (h), haplotype diversity (Hd), nucleotide diversity (Jukes-Cantor), Pi(JC), Tajima D and Fu and Li statistics were performed using obtained TLR3, TLR4 and MHC sequences in the DnaSP6 v6.12.01 (Rozas et al. 2017). The visualization of haplotype diversity for each of the three genes was constructed using TCS haplotype networks in PopArt (Leigh and Bryant 2015). Population pairwise F-statistics for each gene were calculated in Arlequin v3.5, using 10.000 permutations (Excoffier and Lischer 2010). Lastly, the profile of private and shared MHC haplotypes was visualized using the UpSet R package (Conway et al. 2017).

### Selection signatures and recombination analysis

Assessments of the substitution rate and ratio of synonymous and nonsynonymous amino acid substitutions genes across populations were performed using the maximum likelihood YN00 model in PamlX V 1.3.1. (Evolution et al. 2007; Rozas et al. 2017; Xu and Yang 2013). Quantification of diversifying and purifying selection was inferred using the Fast Unconstrained Bayesian Approximation for Inferring Selection (FUBAR) and Single-Likelihood Ancestor Counting (SLAC) methods available on Datamonkey online set of bioinformatic tools (Kosakovsky Pond and Frost 2005; Murrell et al. 2013). GTR substitution model was chosen as the most fitting for MHC and both TLR gene datasets. Selection characters are inferred through the ratio of non-synonymous (d_N_) and synonymous substitutions (d_S_). Positive (diversifying) selection occurs when non-synonymous substitutions are more numerous than synonymous (d_N_/d_S_ > 1). In contrast, the negative (purifying) selection occurs when non-synonymous substitutions are removed from the population due to its deleterious effects, rendering the exon enriched with mostly synonymous mutations (d_N_/d_S_ < 1) (Kryazhimskiy and Plotkin 2008). To analyze whether alleles were more similar at selected sites than at neutral sites, indicating either convergence or trans-species polymorphism (TSP) (Edwards and Hedrick 1998), we performed a comparative analysis using MEGA software. First, nucleotide sequences of MHC class I genes from different bluethroat subspecies were aligned using the ClustalW algorithm in MEGA v7 (Kumar et al. 2016). We then conducted codon-based Z-tests for selection that estimates non-synonymous (d_N_) and synonymous (d_S_) substitution rates using the Nei-Gojobori method with Jukes-Cantor correction implemented in MEGA v7. Phylogenetic trees were constructed separately for non-synonymous and synonymous sites using the Neighbor-Joining method, applying the Jukes-Cantor model with Gamma distribution (+G) and Invariant sites (+I) to account for rate heterogeneity among sites. Average d_N_ and d_S_ values were calculated, and d_N_-d_S_ values were determined for each codon to evaluate selection pressures. The results were interpreted by comparing average d_N_ and d_S_ values: higher average d_N_ would suggest convergence, while higher average d_S_ would suggest TSP. Statistical significance of the d_N_-d_S_ values was assessed using p-values from the Z-tests. To visualize the conserved and variable regions within the peptides and highlight the most frequent amino acids at each position, we generated consensus logo motifs for peptides in three genes (MHC class I exon 3, TLR3, and TLR4) using WebLogo (http://www.weblogo.berkeley.edu/logo.cgi). We grouped peptides for each gene by peptide length and input them into WebLogo with default settings: polar amino acids (G, S, T, Y, C, Q, N) are green, basic (K, R, H) are blue, acidic (D, E) are red, and hydrophobic (A, V, L, I, P, W, F, M) amino acids are black. For each peptide length with sufficient peptides (n > 10), we created a consensus logo motif (Crooks, 2004).

### MHC Supertype inference and analysis of immuno-haemosporidian relationships

To analyze the relationships between MHC class I exon 3 supertypes and haemosporidian infections caused by multiple *Plasmodium* and *Leucocytozoon* lineages we utilized data on prevalence and diversity of haemosporidian fauna in bluethroat populations that were obtained in a previous study (Damnjanović et al. 2025). In short, using species delimitation methods the previous study clustered 19 *Plasmodium* lineages into 11 Operational Taxonomic Units (OTUs), whereas 11 *Leucocytozoon* lineages were found to comprise two OTU units (Damnjanović et al. 2025). Here, we employed GLM analysis (detailed below) to detect associations of functional MHC supertypes with the *Plasmodium* and *Leucocytozoon* OTUs in the bluethroat dataset. Lineages affiliated to the *Haemoproteus* were excluded from analysis due to their very low prevalence (Damnjanović et al. 2025).

The number of MHC supertypes was estimated using the *MHCtools* v1.5.3 R package (Roved et al. 2022). The *DistCalc* function generated five z-descriptor matrices based on fasta format alignment of 1179 MHC class I exon 3 sequences that contained 19 positively selected sites detected by the *FUBAR* analysis (Delport et al. 2010; Kosakovsky Pond and Frost 2005; Murrell et al. 2013). Based on five z-descriptor value matrices, the *BootKmeans* function was executed with a “Hartigan-Wong” algorithm and threshold=0.05 for 1000 number of scans. According to the best AIC value eight putative MHC supertypes were inferred in all 1000 numbers of iterations. The number of the MHC supertypes was determined on an individual level using *adegenet* R package *find.clusters* function with the following settings (max.n.clust=7, n.iter = 1e6) (Jombart 2008). The functional diversity of the MHC class I exon 3 exon proteins was quantitatively expressed through the number of MHC supertypes and qualitatively through presence and absence of each of the eight inferred supertypes separately (Roved et al. 2022; Vlček et al. 2016). The GLM association between MHC class I exon 3 supertypes and prevalence of haemosporidian parasites was performed using regression analyses (generalized linear models in R *lme4* R package) (Bates et al. 2015). Statistical predictors, such as bird subspecies, population, sex, body weight/tarsus ratio, genomic heterozygosity (Damnjanović et al. 2024), number of MHC supertypes and presence/absence of particular supertype were included in order to detect correlation with the infection status with *Plasmodium* and *Leucocytozoon* or mixed case infection (Damnjanović et al. 2025). We designed six regression tests to find correlations between: presence of *Plasmodium* and *Leucocytozoo*n infection status and the total number of MHC supertypes on an individual level (**Regression test 1**); presence of *Plasmodium* and *Leucocytozoon* infection status and the presence or absence of MHC supertypes (**Regression test 2**); quantity of *Plasmodium* and *Leucocytozoon* OTUs detected and the total number of MHC supertypes (**Regression test 3**); quantity of *Plasmodium* and *Leucocytozoon* OTU and the presence or absence of particular MHC supertype (**Regression test 4**); presence of particular *Plasmodium* and *Leucocytozoon* OTU and the total number of MHC supertypes (**Regression test 5**); presence of particular *Plasmodium* and *Leucocytozoon* OTU and the presence or absence of particular MHC supertype (**Regression test 6**).

## Results

### Genetic diversity of the immunity genes

Within three assessed bluethroat populations we recovered 172 unique MHC class I exon 3 alleles. MHC haplotype diversity **Hd** was highest in the Abisko population (0.972 ± 0.004), while similar **Hd** was observed in the Krkonoše (0.966 ± 0.003) and Třeboň (0.962 ± 0.006) respectively. Furthermore, nucleotide diversity (**Pi**) maintained balanced values across populations (Table 1). TLR3 **Hd** showed highly similar values between two red-spotted bluethroat populations (Krkonoše: 0.735 ± 0.026; Abisko: 0.777 ± 0.043, Table 1). The population of white-spotted bluethroat had lower **Hd** (0.623 ± 0.044). Similarly, for TLR4 both red-spotted bluethroat population had highly **Hd** equivalent values (Krkonoše: 0.940 ± 0.007; Abisko: 0.927 ± 0.029), while the white-spotted bluethroat population showed a lower **Hd** score (0.871 ± 0.031). Overall, **the Pi** values of TLR4 gene were higher than TLR3 for all three bluethroat populations (Table 1). Tajima D scores for MHC class I exon 3 had positive non-significant values ranging from lowest (*Tajima D = 0.334*) in the white-spotted bluethroat population in Třeboň area, to the highest (*Tajima D = 0.647*) in Krkonoše population of red-spotted bluethroats. Inversely, Tajima D values for both TLR3 and TLR4 genes had non-significantly negative values across all populations (Table 1).

**Table 1.** Genetic diversity of the immunity genes in three bluethroat populations (**N** - number of samples, **h** - number of haplotypes, **hd** - haplotype diversity, **pi** - mitochondrial nucleotide diversity (Jukes-Cantor), **Tajima D, Fu and Li, K** - number of segregating sites)

| Population | Gene | N | h | Hd (SD) | Pi (JC) | Tajima D | Fu and Li | K |
| --- | --- | --- | --- | --- | --- | --- | --- | --- |
| Krkonoske | MHC | 903 | 156 | 0.966 (0.003) | 0.0792 | 0.64691 | -24.691 | 137 |
|  | TLR3 | 243 | 21 | 0.735 (0.026) | 0.0041 | -1.52038 | -12.382 | 20 |
|  | TLR4 | 283 | 63 | 0.940 (0.007) | 0.0059 | -1.50826 | -33.964 | 34 |
| Abisko | MHC | 169 | 65 | 0.972 (0.004) | 0.0785 | 0.39978 | -3.588 | 116 |
|  | TLR3 | 42 | 10 | 0.777 (0.043) | 0.0035 | -1.43795 | -4.855 | 10 |
|  | TLR4 | 41 | 23 | 0.927 (0.029) | 0.0052 | -1.77444 | -22.140 | 20 |
| Třeboň | MHC | 107 | 38 | 0.962 (0.006) | 0.0767 | 0.33353 | 2.041 | 106 |
|  | TLR3 | 41 | 5 | 0.623 (0.044) | 0.0022 | -0.46632 | -1.071 | 4 |
|  | TLR4 | 45 | 14 | 0.871 (0.031) | 0.0056 | -0.81851 | -5.135 | 14 |

The majority of TLR3 and TLR4 haplotypes differ only in one nonsynonymous polymorphic site (Fig. 2). The large number of mutually interconnected TLR4 haplotypes are present due to a high number of synonymous substitutions, while TLR3 displayed less synonymous mutations and haplotype diversity. The haplotype network of MHC class I exon 3 was increasingly more complex, in which irregular constellations across populations to did not allow any clear resolution. Contrastingly, translated haplotype network showed distinction between eight amino acid sequences (Supplementary Material. Fig. S1.) Population pairwise F_ST_ analysis revealed a small but statistically significant (*p < 0.05*) population structuring in TLR 3 genes between the two red-spotted bluethroat populations (*F*_*ST*_ *=0.02248*). Additionally, a significant difference in the structure of TLR4 genes was found between Krkonoše an Třeboň populations (*F*_*ST*_ *= 0.02078*).

**Fig. 2.**
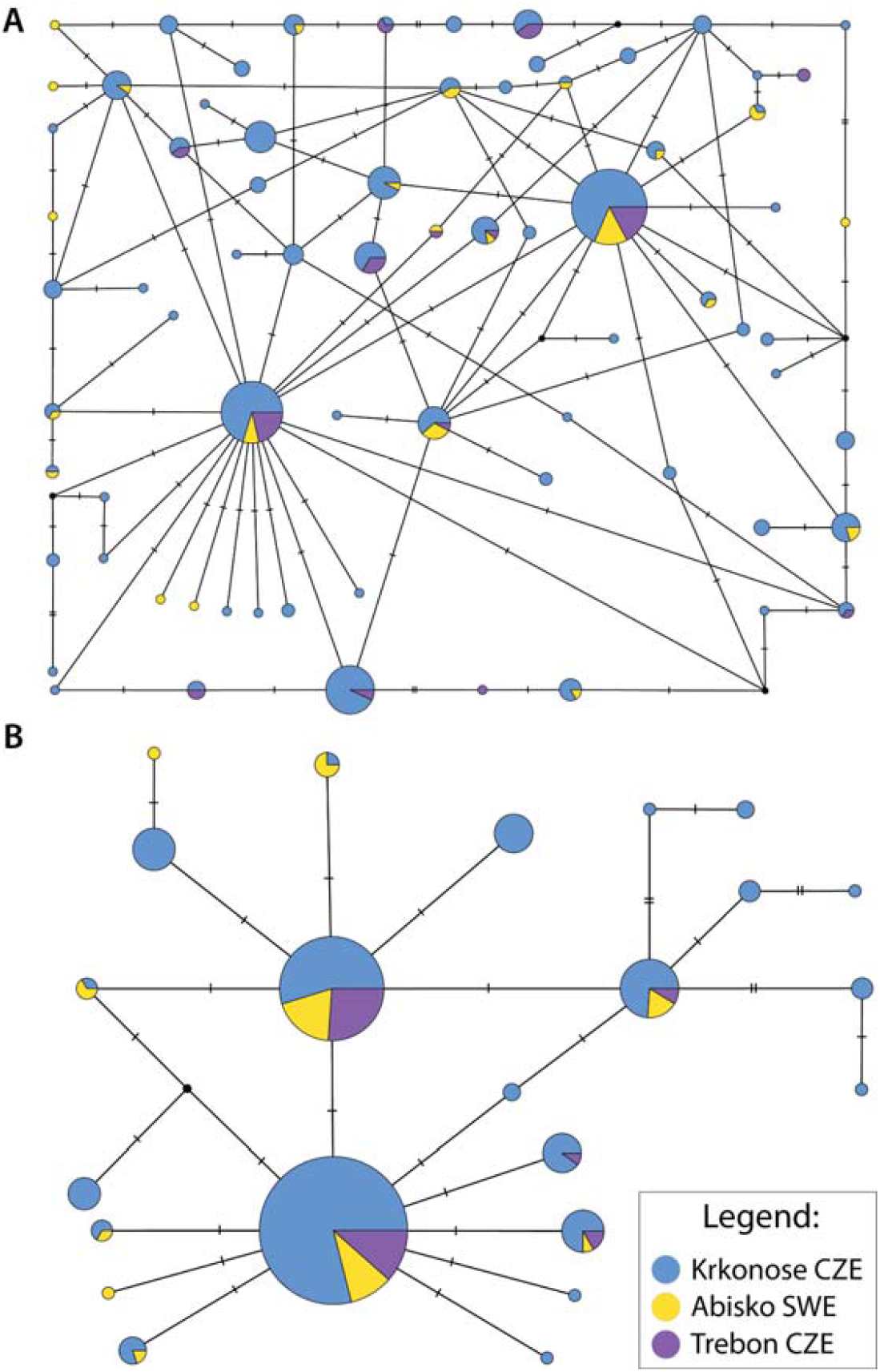
TCS haplotype network depicting abundance and diversity of TLR4 (A) and TLR3 (B) haplotypes across populations. Several mutational steps between haplotypes are depicted by hatch marks.

The genetic diversity analysis of MHC haplotypes identified 10 common haplotypes across all three populations. Reflecting its largest sample size, the bluethroat population in Krkonoše exhibited the highest diversity, with 156 haplotypes, in contrast to Třeboň, which had the lowest number of unique haplotypes (38 haplotypes). Furthermore, the populations from Krkonoše and Abisko shared the most haplotypes (49 haplotypes). I comparison, Krkonoše and Třeboň shared 18 haplotypes, and Abisko and Třeboň shared 6 (Fig. 3). The results of pairwise F_ST_ demonstrated that MHC genes of white-spotted bluethroats in Třeboň significantly (*p < 0.05*) diverged from both red-spotted populations located in Abisko (*F*_*st*_ *= 1.708*) and Krkonoše (*F*_*st*_ *= 0.971*). However, the genetic structure of MHC genes between Abisko and Krkonoše did not show any significant difference (*F*_*st*_ *=0.007*).

**Fig. 3.**
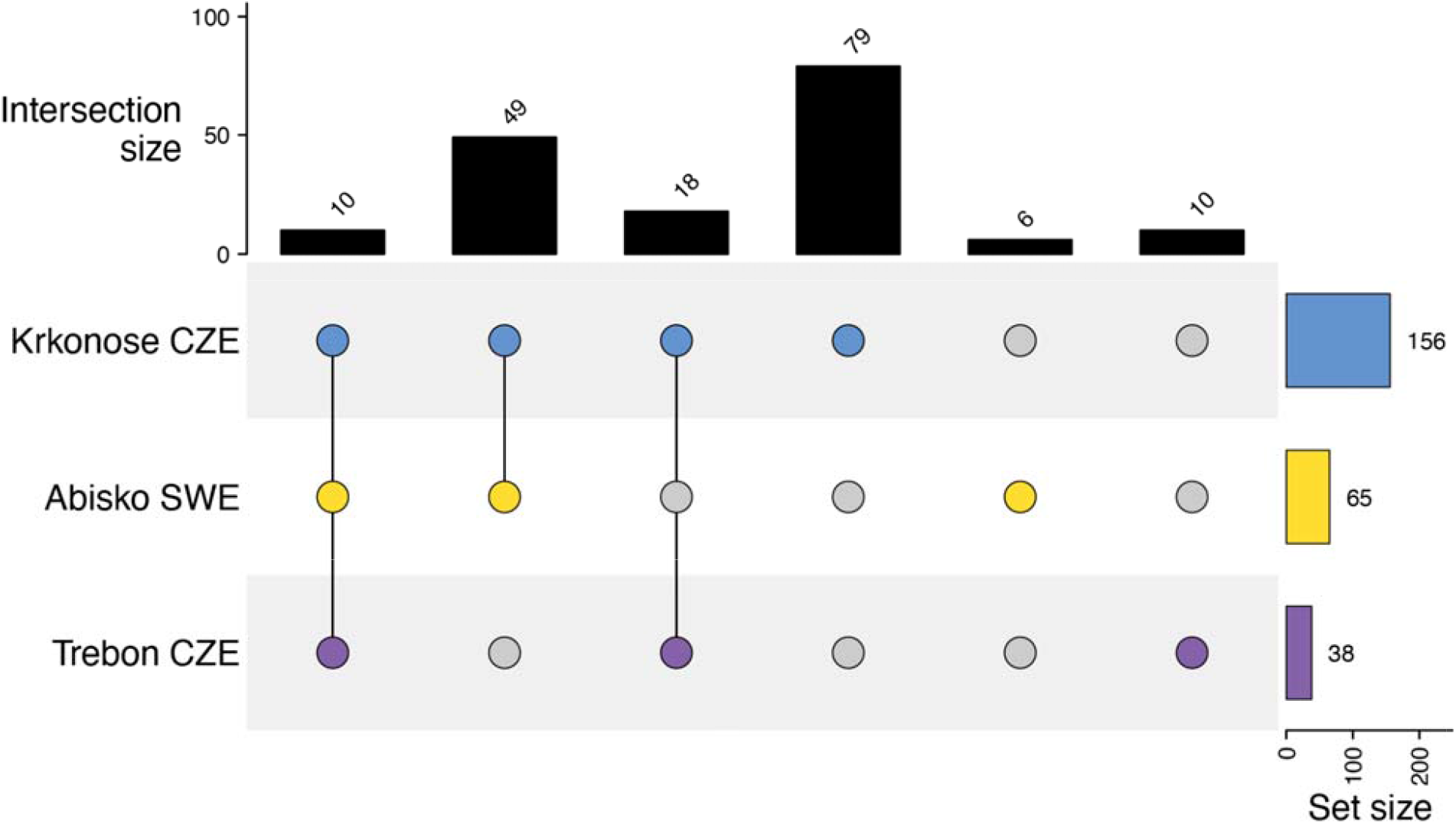
Profile of specific and shared MHC alleles between populations. The horizontal axis of the UpSet plot represents the number of MHC nucleotide haplotypes shared between populations and the number of populatio specific haplotypes. The vertical axis depicts the total number of MHC haplotypes per population.

### Selection analysis of the immunity genes

TLR3 gene in bluethroat species possesses only a few amino acid sites affected by diversifying and purifyin selection (Table 2). Namely, the *FUBAR* method identified a single site at the 6th position under positive selection and two sites at positions 112 and 118 that are negatively selected. In contrast, the *SLAC* method only detected a single amino acid site at position 118, showing negative selection (*AICc = 1404.08*). Krkonoše population containe a single amino acid at the site 6 favored by positive selection, while white-spotted bluethroat in Třeboň showed the sites 85 and 102 under positive selection (Table S1). Conversely, purifying selection in TLR4 gene was notably predominant. The *FUBAR* and *SLAC* analysis did not find any signs of diversifying selection within the TLR4 gene, whereas ten amino acid positions (*site no: 1, 35, 45, 64, 70, 90, 97, 105, 108, 125*) were found under the effect of purifying selection by both methods (*AICc - 2644.04*) (Table 2). On the population level, only the amino acid sites 90 and 125 were negatively selected in all three bluethroat populations, while other sites were population specific (see Table S1).

**Table 2.** Summary of selection analysis for triplet of immunity genes in bluethroat performed by FUBAR, SLAC and PAML analysis; **UnH/UnAA** - unique number of haplotypes / unique number of amino acid variants).

| Exon | N selection sites<br>FUBAR |  | N selection sites<br>SLAC |  | dN/dS ratio<br>(YN00) | Z test substitution<br>rates | UnH/UnAA |
| --- | --- | --- | --- | --- | --- | --- | --- |
|  | positive | negative | Positive | negative |  |  |  |
| MHC | 19 | 20 | 13 | 16 | 0.8430 |  | 172/137 |
| TLR3 | 1 | 2 | 0 | 1 | 0.0854 |  | 18/10 |
| TLR4 | 0 | 10 | 0 | 10 | 0.0873 |  | 69/17 |

In contrast to both TLR genes, the MHC class I exon 3 gene displays a dense arrangement of amino acid sites that are simultaneously under the effect of positive and negative selection. Namely, *FUBAR* identified 19 (*8, 10, 13, 23, 26, 32, 35, 41, 58, 61, 64, 75, 80, 85, 93, 100, 102, 108, 110*) and 20 (*7, 18, 19, 20, 21, 29, 30, 31, 39, 40, 43, 45, 48, 69, 76, 81, 83, 84, 90, 106*) amino acid sites that are under effect of positive and negative selection, respectively (Table 2). Similar results were obtained from the SLAC analysis where 13 amino acid sites under positive selection were identified (*8, 10, 26, 32, 41, 58, 64, 75, 80, 85, 93, 100, 108*) and 16 under negative selection (*6, 7, 19, 20, 21, 29, 30, 31, 40, 43, 45, 48, 63, 69, 83, 84*). The selection analysis performed by the YN00 model inferred the presence of negative selection affecting the evolution within all three immunity genes (*MHC: d*_*N*_*/d*_*S*_ *= 0.8430; TLR3 d*_*N*_*/d*_*S*_ *= 0.0854; TLR4 d*_*N*_*/d*_*S*_ *= 0.0873*). All three bluethroat populations showed positive and negative selection on an identical collection of amino acid sites (*10, 26, 32, 58, 64, 75, 100, 108*), while other MHC class I exon 3 amino acid sites have a population specific character (Table S1.). Conversely ten negatively selected sites that were found in all three assessed populations (*7, 19, 30, 31, 43, 45, 48, 69, 83, 84*).

The Z test substation rate inferred that the number of d_S_ and d_N_ between two red-spotted bluethroat populations was similar (d_S_ = 20; d_N_ = 17). Similar amount was observed also between the Krkonoše and Třeboň population (d_S_ = 21; d_N_ = 14). Additionally, the results of codon-based Z test for selection and subsequent phylogenetic analysis of MHC gene revealed that the average non-synonymous substitution rate (d_N_) was 2.135, while the average synonymous substitution rate (d_S_) was 4.289. Moreover, the average difference between non-synonymous and synonymous substitution rates (d_N_-d_S_) was −2.154. Moreover, the normalized average d_N_-d_S_ value was −0.94. Similarly, the d_N_ and d_S_ values for of Z test for both TLR3 (d_S_ = 0.19; d_N_ = 0.06;) and TLR4 (d_S_ = 0.69; d_N_ = 0.006). Namely, Multiple TLR3 and TLR4 haplotypes are found within both bluethroat subspecies (Fig 2.).

The sequence logo plots for three genes, MHC class I, TLR3, and TLR4 highlight distinct patterns of variability across the peptide sequences (Fig. 4). For MHC class I, the sequence logo revealed significant variability at most amino acid positions (*e.g*., *10, 26, 28, 32, 33, 58, 64, 85, 108*), indicating a high degree of diversity in this gene. This variability suggests that the MHC class I exon 3 protein can present a wide range of peptides, which is critical for the immune system’s ability to recognize diverse pathogens (Fig. 4A). However, the TLR3 gene shows moderate conservation with notable variability at positions such as 6, 40, 51, and 81 (Fig. 4B). The variability in these positions might reflect the receptor’s ability to recognize various pathogen-associated molecular patterns. For TLR4, the sequence logo displays a mix of highly conserved and highly variable positions. Positions such as 28 and 34 are particularly variable in these codon positions (Fig. 4C).

**Fig. 4.**
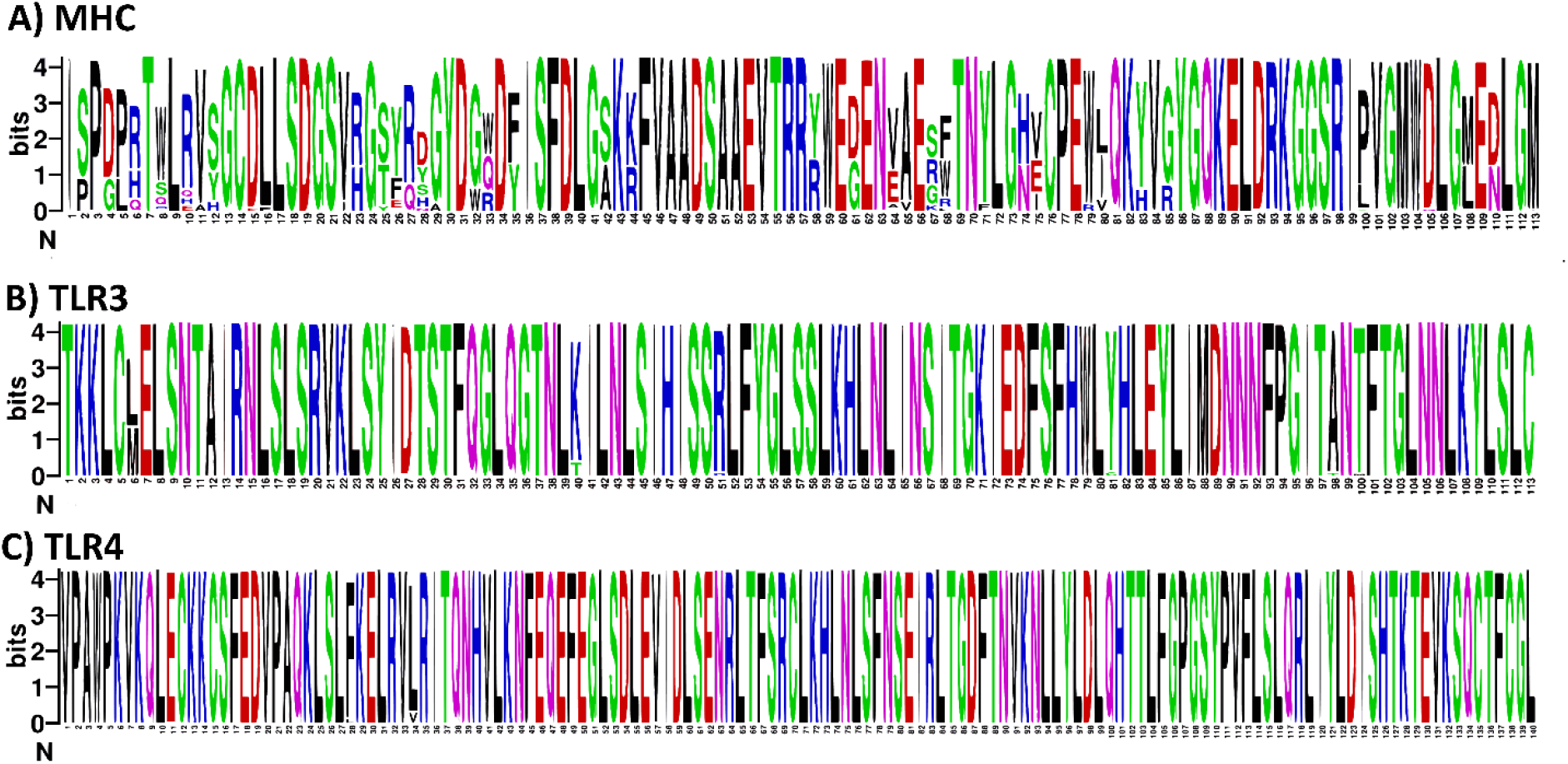
Sequence Logo plots representing amino acid diversity of the three immune genes in bluethroats: (A) Major Histocompatibility Complex (MHC), (B) Toll-Like Receptor 3 (TLR3), and (C) Toll-Like Receptor 4 (TLR4). Each logo illustrates the nucleotide or amino acid frequencies at each position, with the height of the letters indicating the relative frequency of each base or residue. Different colors represent different types of amino acids, providing a visual representation of the sequence variability and conservation within these genes.

### Supertype relationship with the presence of the haemosporidian infection

No statistically significant relationship was found between general presence of *Plasmodium, Leucocytozoon* or mixed infections with the number of MHC supertypes per individual (Fig. 5) (**regression test 1**). The presence or absence of any MHC supertype was not linked with the overall infection status with *Plasmodium* or *Leucocytozoon* genus (**regression test 2**). Also, the number of detected *Plasmodium* and *Leucocytozoon* OTU units was not linked with the number of MHC supertypes per individual (**regression test 3**). No significant link was found between the total number of detected *Plasmodium* and *Leucocytozoon* OTU and the presence and absence of the eight MHC supertypes on an individual level (**regression test 4**). None of the putative haemosporidian OTU units was linked with the total number of MHC supertypes (**regression test 5**). Lastly, **regression test 6** detected three statistically significant correlations. Firstly, we detected a statistically significant positive association between *Plasmodium* OTU3 and MHC supertype 6 (*p = 0.009; 𝒳2 = 1.95*). Second correlation was detected between *Plasmodium* OTU5 and MHC supertype 3 (*p = 0.01; 𝒳2 = −27.75*). Lastly, *Leucocytozoon* OTU2 have simultaneously a positive association with MHC supertype 4 (*p = 0.04;* χ*2 = 3.732e + 01*) and a negative with the MHC supertype 6 (*p = 0.01;* χ*2 = 3.682e+01*).

**Fig. 5.**
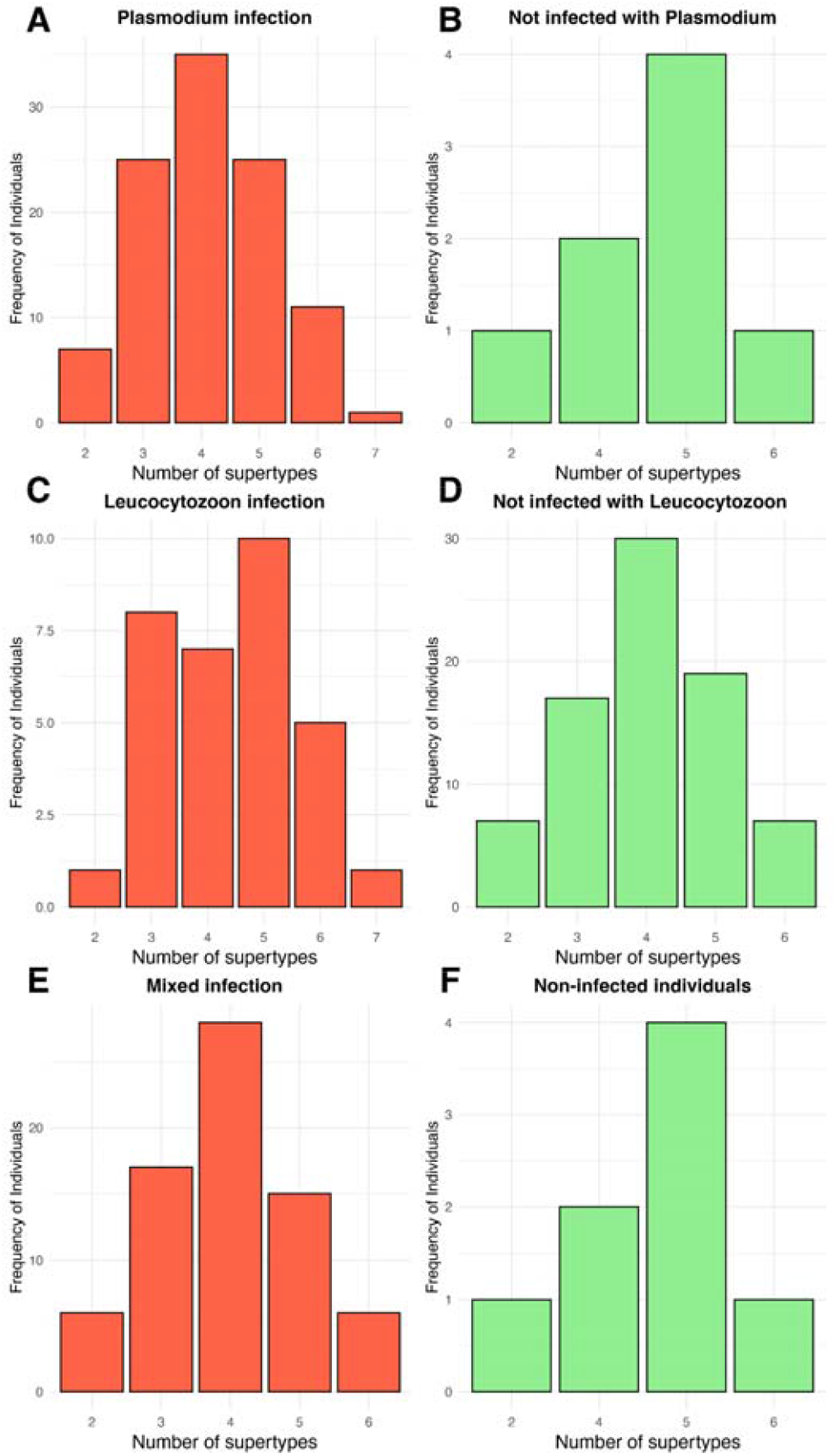
Numbers of MHC supertypes between infected (red) and noninfected (green) bluethroats in *Plasmodium* (A-B), *Leucocytozoon* (C-D) and mixed case infections (E-F).

## Discussion

### Genetic diversity of immunity genes

In this study we reported the genetic diversity of three immunity genes (TLR3, TLR4 and MHC I exon 3) within the three European bluethroat populations. We also carried out selection analysis by *FUBAR* and *SLAC* methods that provided compact selection profiles acting on focal immune genes. Lastly, while the genetic diversity in TLR genes was low for testing correlations between particular gene variants and haemosporidian infection status, it was performed using GLM regression analysis for MHC gene variants and haemosporidian parasites through six regression tests. In combination with the DADA2 R package, a triplet of in-house modified primer pairs performed with great accuracy in retrieving high quality sequences from all three assessed genes (Alcaide et al. 2013; Callahan et al. 2016; Lyons et al. 2015). We have documented a great similarity in nucleotide and haplotype diversity across all three bluethroat populations despite their geographical and ecological differences and varying population size. More importantly, we found that most frequent haplotypes were present in both the red-spotted (Krkonoše and Abisko) and white-spotted (Třeboň) bluethroat subspecies (Fig. 2; Supplementary Material, Fig. S1). Low positive Tajima D values in MHC class I exon 3 genes across all three populations potentially pinpoints a slight decrease of genetic polymorphism over time. Conversely, negative Tajima D values in all TLR genes depict a mild increase in both low frequency polymorphisms (Table 1).

The Krkonoše population of red-spotted bluethroats is relatively young as the first individuals were recorded in 1970’s, probably established as migrants from northern Europe (Miles and Formánek 1989). In comparison to the main breeding range in boreal Eurasia, the remote Krkonoše population is considered to have an insular biogeographical character of red-spotted bluethroat subspecies (Macarthur and Wilson 2021). The population was relatively stable in the first decade since colonization, which was, however, followed with a gradual decline over the years. (Miles and Formánek, 1989; Chutný, 1991). Currently, the extant population counts no more than a few singing males recorded in the last decade (personal observation). Surprisingly, although the red-spotted bluethroat population in Krkonoše is at the brink of extinction and possibly permeated with inbreeding load from past consanguineous mattings, it does not show a significantly reduced genetic diversity in any of the three focal immune genes (Table 1. Correspondingly, also Damnjanović et al. 2024) found only a moderate level of diversity reduction using whole genome SNP analysis. Similarly, genetic diversity of TLR genes in insular avian populations can be affected by genetic drift (e.g. Vlček et al., 2022), genetic diversity statistics of immunity genes in the red-spotted bluethroat Krkonoše population refutes its “insular” character.

### Selection analysis of TLR genes revealed dominance of purifying selection

The three immunity genes (TLR3, TLR4 and MHC class I exon 3) were analyzed for signatures of positive and negative selection. High contrast of evolutionary forces that shape TLR innate genes in comparison to the MHC adaptive immunity genes has been observed by the different ratio of amino acid sites under positive and negative selection (Fig. 4, Supplementary Material Table S1). Notably, the higher presence of purifying selection pinpoints a higher ratio of synonymous mutations over the non-synonymous, whereas minor presence of the positive selection introduced few nonsynonymous substitutions. Similar character of selection pressures on TLR genes was reported in a wide range of avian taxa, pinpointing conserved function of innate immunity genes (Grueber et al. 2014; Nelson□Flower et al. 2023; Velová et al. 2018; Yilmaz et al. 2005). The constrained character of the TLR gene evolution is associated with the archaic structure and highly conserved function of the TLR genes. Predominant purifying selection is characterized by low allelic richness allowing predominantly synonymous polymorphism (Boyd et al. 2018; Chapman et al. 2016; Grueber et al. 2012). Specifically, out of the 69 unique haplotypes identified within the whole bluethroat TLR4 dataset, we detected 48 synonymous nucleotide haplotypes that translated into a single amino acid haplotype. Both TLR3 and TLR4 exhibit low d_N_/d_S_ values that signify strong purifying selection, implying that changes to these genes are predominantly deleterious and are being selectively removed (Yang et al. 2000, 2005). Similarly, other inborn immune genes in avian species such as ß-Defensins have shown low diversity on both individual and population level (Chapman et al. 2016). Consequently, due to their essential role and conserved nature of inborn immune genes as a whole, non-synonymous substitutions are a profound change that abate its functionality (Roach et al. 2005). Conversely, scarce signatures of positive selection affecting evolution have been reported in a wide range of vertebrate taxa (Boyd et al. 2018; Grueber et al. 2012, 2014; Roach et al. 2005).

### MHC Class I exon 3 is governed by balanced influence of diversifying and purifying selection

Genetic analysis revealed the presence of both diversifying and purifying selection forces that synchronically affect different amino acid sites of the MHC class I exon 3 sequences in bluethroat species (Table 2, Supplementary Material Table S1). Multiple amino acid sites of the MHC class I exon 3 gene is mutually governed by two opposing selection directions that consequently generate a combined effect on the maintenance of diversity of the MHC genes (Antonides et al. 2017; Spurgin and Richardson 2010). The presumable effect of balancing selection could be at work in three populations due to great similarities in MHC genetic diversity (Cruz-López et al. 2020). Our results depicts that selection affecting MHC class I exon 3 gene remain in equilibrium across populations, despite the population size, haplotype diversity, population structure, and diversity of local haemosporidian fauna (Damnjanović et al. 2025; Damnjanović et al. 2024; Svoboda et al. 2015a). The analysis of d_N_/d_S_ ratios for three genes revealed presence of selective pressure that maintains variation within MHC class I exon 3 gene. Mixing of MHC alleles within and across breeding pairs might be shaped in a non-random manner (Milinski 2006; Rekdal et al. 2019; Woelfing et al. 2009). Interestingly, to increase fitness of offspring survival, female bluethroats/passerines presumably choose their male mate that have complementary allelic diversity that will set up optimal MHC array of alleles. Specifically, in female bluethroats it was reported that female alter their mating preferences in accordance with availability of specific age-category males (Rekdal et al. 2023), which in turn can affect the dispersal of MHC alleles within and across populations.

According to the trans-species evolutionary model, MHC orthologs are inherited from ancestral species to two or more descendant species or subspecies (Agudo et al. 2012; Edwards and Hedrick 1998). Our results demonstrate that four out of eight MHC supertypes are found in both bluethroat subspecies (Fig. 3, Supplementary material, Fig. S1). Conclusively, balancing selection results in preserving the ortholog alleles across multiple subspecies of species (Cutrera and Lacey 2007; Kasahara et al. 1995). The average d_S_ value (4.289) is higher than the average d_N_ value (2.135), indicating that synonymous (neutral) sites have higher average substitution rates compared to non-synonymous (selected) sites, advocating stronger conservation at non-synonymous sites. Classical TSP hypothesis states that ancestral alleles are maintained and show higher similarity at neutral sites. Moreover, the negative value of the average d_N_-d_S_ (−2.154) indicates that synonymous substitutions are more frequent than non-synonymous ones, supporting the idea that non-synonymous sites are under stronger purifying selection, consistent with TSP (Eimes et al. 2015, 2016; Klein et al. 1998). Given that prevalence and diversity of haemosporidian fauna in the same populations have been reported to show a significant difference in the beta diversity for the *Plasmodium* and *Leucocytozoon* lineages (Damnjanović et al. 2025), local haemosporidian fauna might impose a minute impact on the selection of the MHC class I exon 3 gene on a population level (Table 1).

### MHC class I exon 3 relationship with *Plasmodium* and *Leucocytozoon* infections

We used MHC class I exon 3 supertypes as representative units of quantitative and qualitative diversity on individual level to test general linear regression models against the presence of various lineages of *Plasmodium* and *Leucocytozoon* grouped into OTUs defined by Damnjanović et al. (2025). We report that haemosporidian (*Plasmodium* and *Leucocytozoon*) infection status and number of haemosporidian OTU carried per are not predicted by sex nor morphometric indices. We also report that high MHC allelic heterozygosity was not linked with lower prevalence or absence of infection than in individuals with a lower number of MHC supertypes. Individuals with both high and low number of MHC supertypes had equal probability of being infected with both *Plasmodium* and *Leucocytozoon* parasites (Fig. 5). Secondly, quantity of the MHC alleles itself does not necessarily provide a cumulatively stronger immunological response against each type of a pathogen (Jones et al., 2015; Sutton et al., 2016; Worley et al., 2010). None of the MHC supertypes with low frequency were correlated with any of the delimited *Plasmodium* or *Leucocytozoon* OTU units. Our findings did not provide support for the *heterozygote advantage* nor *rare allele advantage* hypothesis in the relationship between bluethroat MHC genes and diversity of reported haemosporidian fauna in bluethroat species (Edwards and Hedrick 1998; Spurgin and Richardson 2010).

The sole presence or absence of the particular MHC supertype can have a significant contribution to the outcome of infection and parasitemia (Nelson□Flower et al., 2023; Vlček and Štefka, 2020; Westerdahl et al., 2012). In this study, we have discovered a few positive and negative correlations of specific MHC supertypes with the presence of a specific haemosporidian OTU unit (**Regression test 6**). Several studies have reported negative associations of certain MHC supertypes with the presence of parasites. It is assumed that these correlations impose a qualitative immunity, rendering host individuals’ infection free (Biedrzycka et al. 2018; Sepil et al. 2013). In contrast, positive associations between MHC supertypes and parasite infections provides quantitative resistance, where reduced parasitemia impose minimal disadvantage on individual fitness and daily dynamics (Vlček and Štefka 2020; Westerdahl et al. 2012). Moreover, variability of environmental factors and geographical features can further alter the probability of infection and spatial distribution of parasites (Jones et al. 2015; Lazzaro and Little 2009; Mitchell et al. 2005). Several studies reported a detrimental effect of specific MHC supertypes on longevity of individuals (e.g., Nelson□Flower et al., 2023). Presumably, a study that tests multiple types of pathogens (i.e. viral, bacterial, fungal and malarial) would deem a more plausible framework in which a *heterozygous advantage* model of balancing selection could be more accurately tested.

## Acknowledgements

We thank several colleagues, namely Jakub Vlček and Helena Westerdahl, for discussing details of some laboratory and data analyses with us. Computational resources were provided by the e-INFRA CZ project (ID:90254), supported by the Ministry of Education, Youth and Sports of the Czech Republic.

## Authorship contribution

Conceptualization: Dragomir Damnjanović, Masoud Nazarizadeh, Jan Štefka; Writing – original draft: Dragomir Damnjanović, Masoud Nazarizadeh, Jan Štefka; Sample acquisition: Václav Pavel, Arild Johnsen, Bohumír Chutný; Methodology and Formal analysis: Dragomir Damnjanović, Masoud Nazarizadeh, Milena Nováková, Jan Štefka; Supervision and Funding acquisition: Jan Štefka. All authors read and approved the final manuscript.

## Data availability

All NGS amplicon sequencing data generated in this study are available in the National Center for Biotechnology Information (NCBI) Sequence Read Archive under BioProject accession PRJNA1265452.

