## Supplementary figures and images for "Immunogenetic diversity and haemosporidian parasitization in European bluethroats (*Luscinia svecica*): do diverse genes warrant fewer parasites?"

### Supplementary Fig. S1

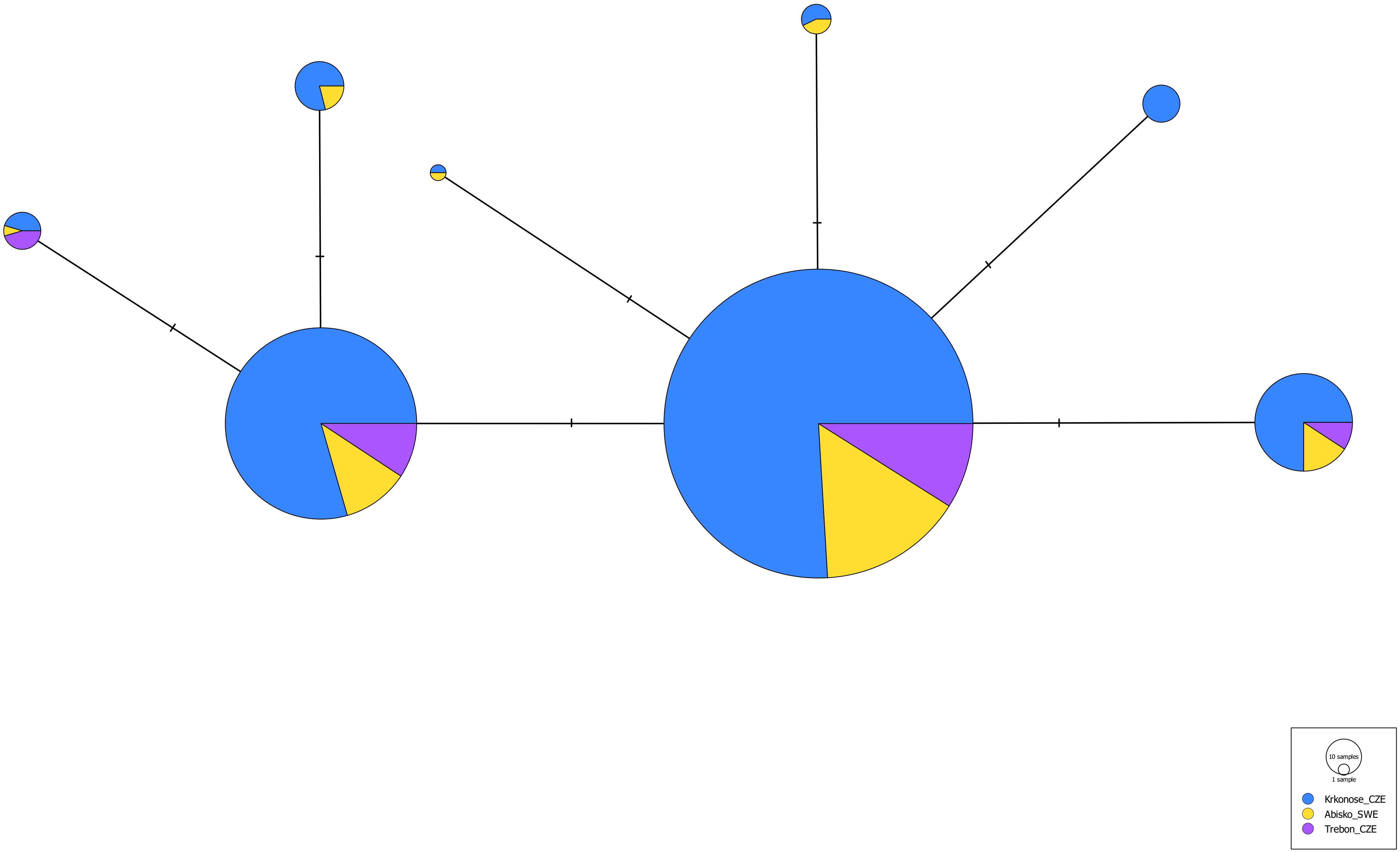
